# Dissecting the molecular puzzle of the editosome core in Arabidopsis organelles

**DOI:** 10.1101/2024.01.16.575875

**Authors:** Kevin Baudry, Dario Monachello, Benoît Castandet, Wojciech Majeran, Claire Lurin

## Abstract

Over the last decade, the composition of the C-to-U RNA editing complex in embryophyte organelles has turned out to be much more complex than first expected. While PPR proteins were initially thought to act alone, significant evidences have clearly depicted a sophisticated mechanism with numerous protein-protein interaction involving PPR and non-PPR proteins. Moreover, the identification of specific functional partnership between PPRs also suggests that, in addition to the highly specific PPRs directly involved in the RNA target recognition, non-RNA-specific ones are required. Although some of them, such as DYW1 and DYW2, were shown to be the catalytic domains of the editing complex, the molecular function of others, such as NUWA, remains elusive. It was suggested that they might stabilize the complex by acting as a scaffold. We here performed functional complementation of the *crr28-2* mutant with truncated CRR28 proteins mimicking PPR without the catalytic domain and show that they exhibit a specific dependency to one of the catalytic proteins DYW1 or DYW2. Moreover, we also characterized the role of the PPR NUWA in the editing reaction and show that it likely acts as a scaffolding factor. NUWA is no longer required for efficient editing of the CLB19 editing sites once this RNA specific PPR is fused to the DYW catalytic domain of its partner DYW2. Altogether, our results strongly support a flexible, evolutive and resilient editing complex in which RNA binding activity, editing activity and stabilization/scaffolding function can be provided by one or more PPRs.

## INTRODUCTION

In embryophyte organellar transcripts, RNA editing specifically deaminates hundreds of cytidines into uridines. It is catalyzed by a protein complex called “editosome” that contains several members of various nuclear encoded protein families such as Pentatrico Peptide Repeat (PPR), Multiple Organellar RNA editing Factor / RNA-editing factor Interacting Protein (MORF/RIP) or Organelle RNA Recognition Motif-containing (ORRM) proteins (1–4). Among all these editing factors, the PPR proteins were shown to be the *trans* specificity factors that bind the RNA *cis* recognition elements, therefore allowing the specific targeting of the edited cytidines (5, 6).

PPR proteins are part of the ‘α-solenoid’ superfamily and are hypothesized to derive from Tetratrico Peptide Repeat (TPR) proteins (7, 8). They are characterized by repetitions of a degenerate 35 amino acids motif called the PPR motif. The PPR proteins can be classified in two groups based on the nature of their PPR motifs. When only canonic motifs (P motifs) are present, the proteins are called pure PPR defining the P subfamily. When motif variants are observed together with P motifs, the proteins are called PLS PPR proteins and the motifs are called “S motif” for the smaller ones and “L motifs” for the longer ones. PPR proteins can specifically bind RNA in a 1 motif – 1 nucleotide interaction. The binding specificity is explained by a probabilistic degenerate code based on the 5^th^ and 35^th^ amino acid couple of each motif (9–11). Almost all PPR proteins involved in RNA editing are members of the PLS subgroup and harbor additional C-terminal domains named E1, E2, E+ and DYW domains (12, 13) (Fig. 1A).

**FIGURE 1.**
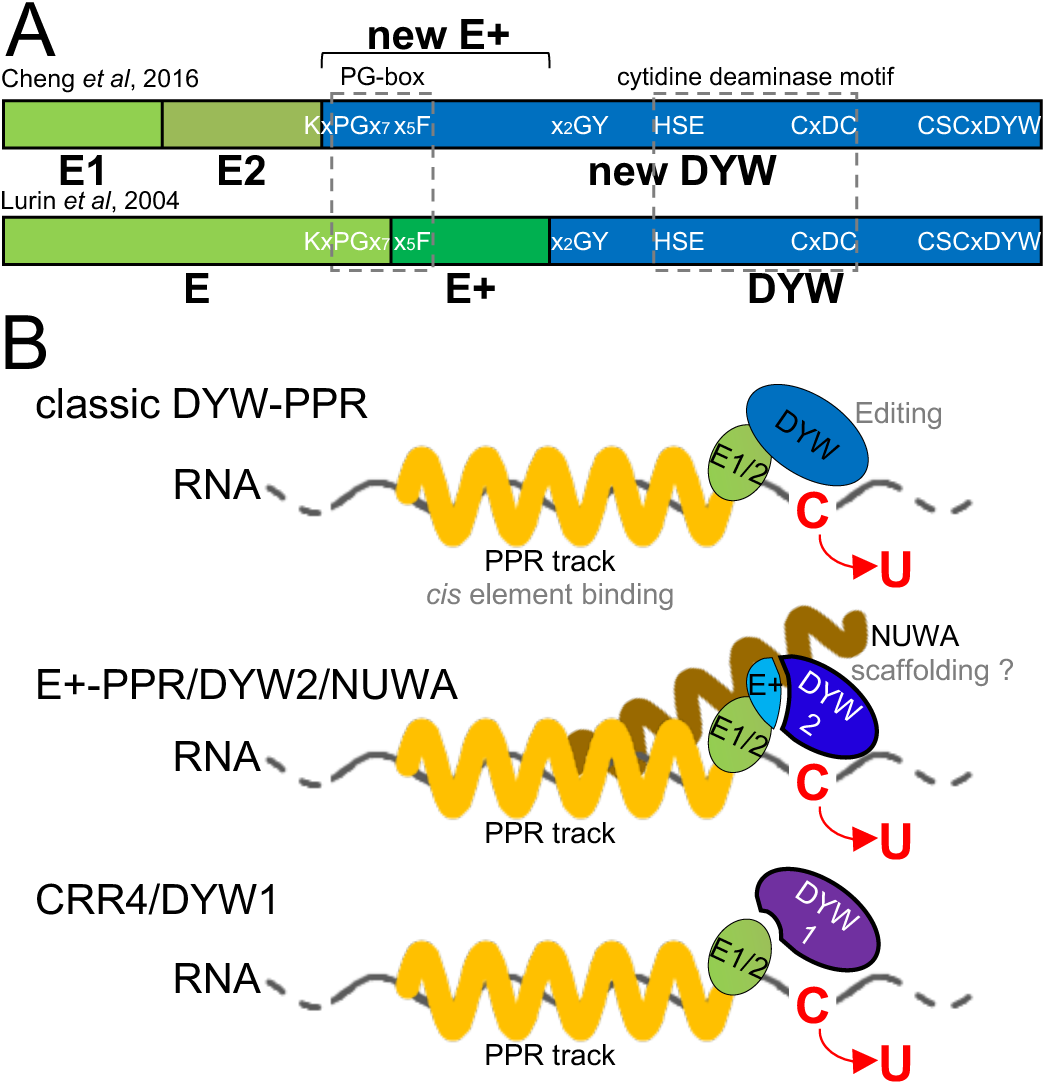
C-terminus domains of editing PPR and chloroplast core editosome models. A) Comparison of C-terminus domain boundaries between Lurin *et al.* (2004) and Cheng *et al.* (2016). In the present work, the new E+ domain is defined as the N-terminus region of new DYW (from Chen et al., 2016) ending where old DYW (from Lurin et al., 2004) starts. B) In chloroplast, three editosome cores were described depending on the subfamily of the site specific PPR involved. Top, DYW-PPR proteins carry the domains required for site binding and editing in the same protein. Middle, an E+-PPR protein binds the site that is edited by a second PPR protein, DYW2. These two PPRs are bridged by a third PPR, NUWA. Bottom, CRR4 (E2-PPR) binds the site and recruits a second PPR, DYW1, to edit it.

Although the identity of the enzyme responsible for the deamination reaction has long remained elusive, it is now clear that the DYW domains at the C-terminus of some PPR proteins carry the catalytic activity. It was for example shown that two *Physcomitrium patens* DYW-PPRs (PpPPR56 and PpPPR71) truncated of their DYW domain (ΔDYW-PPR) are unable to complement their respective mutants (14). Moreover, the DYW-PPR PpPPR65 has been shown to be sufficient to drive editing of its cognate editing site in both *in vitro* assays and *in vivo* heterologous *E. coli* system (15, 16). Moreover, point mutation within any of the DYW conserved deaminase signature motif and zinc binding domains strongly inhibits or abolishes *in vivo* editing efficiency (16–22). It also contains additional domains like the “PG-box”, a 15 amino acid motif encompassing the E and E+ domains according to the domain definition from Lurin *et al*, 2004 (12), and is now included at the beginning of the DYW domain according to more recent definition (13) (Fig. 1A). Finally, structural analyzes of the DYW domain of the plastid RNA editing factor ORGANELLE TRANSCRIPT PROCESSING 86 (OTP86) found it to be highly similar to those of classical cytidine deaminases with some additional specific features. It notably contains a gating domain between the PG-box and the cytidine deaminase motif that is likely involved in the regulation of the active site (23).

Different from *P. patens*, many ΔDYW-PPRs truncated proteins are however able to complement their respective mutants in *Arabidopsis*, suggesting a non-essential role of the DYW domain (20, 22, 24, 25). This result is in agreement with the identification of numerous PPR editing factors that lack the DYW domain and belong to the E2-or E+-PPR subfamily, such as CHLORORESPIRATORY REDUCTION 4 (CRR4), a plastidial E2-PPR or CHLOROPLAST BIOGENESIS 19 (CLB19) and SLOW GROWTH 2 (SLO2), respectively plastidial and mitochondrial E+-PPR proteins in *Arabidopsis thaliana* (5, 26, 27). The apparent dispensability of the DYW domain in flowering plants remained a conundrum until the identification of a small group of atypical short DYW-PPRs, named DYW1-like proteins after DYW1, the first protein of the subfamily to be identified (28). In *Arabidopsis*, the six members of this group differ from classic DYW-PPR proteins by harboring very few PPR motifs (from 0 to 6) and non-canonic E1, E2 and E+ domains (28–30). This led to a model in which each truncated PPR protein subfamily specifically recruits one of the members of the DYW1-like subfamily in order to reconstitute an active PPR-DYW with all the necessary domains (28, 30–32) (Fig. 1B). It was shown for example that DYW1 interacts with CRR4 to edit the plastidial ndhD_117166 site (28), that all editing sites known to be depending on an E+-PPR also depend on DYW2 (30, 33–35) and that MEF8/8S are recruited by all E2-PPR in mitochondria (36).

Finally, many additional, non PLS-PPR proteins, are known to be required for RNA editing, like the MORF proteins on one hand, and NUWA, DG409 and GRP23, three pure PPR proteins, on the other hand (2, 30, 36–38). NUWA is hypothesized to be involved in the stabilization of the interaction between E+-PPR and DYW2 in maize and *Arabidopsis* and GRP23 in the stabilization of the interaction between E2-PPR, DYW-PPR and MORF proteins (30, 33, 39, 40). All these interactions are essential to RNA editing in flowering plants.

Here, we systematically tested the hypothesis that the interaction between DYW1-like proteins and E2/E+-PPR proteins reconstitutes an editing factor with both RNA binding and deaminase activities and that this interaction is supported by scaffolding proteins. We first show that the essential function of the PPR NUWA can be ascribed to its role in scaffolding the interaction between the RNA binding PPR CLB19 and the deaminating PPR DYW2. We then show that functional complementation of the *crr28-2* mutant (a plastidial DYW PPR) with truncated proteins mimicking either an E+- or E2-PPR are fully explained by their respective specific dependencies to DYW2 or DYW1. Our results strongly support a flexible and resilient editosome model, in which the binding of the *cis* recognition element and the cytidine deamination can be carried by one or several PPR as long as all the essential PPR editing domains (PPR tracks and DYW) are present in the editosome core.

## RESULTS

### Plants expressing a CLB19-DYW2 fusion do not need NUWA to edit the CLB19 dependent editing sites

NUWA is believed to be a scaffolding protein that could specifically stabilize E+-PPR/DYW2 complexes. To test this hypothesis, we fused the DYW domain of DYW2 to the C-terminus of the E+ domain of CLB19, a plastidial E+-PPR that requires NUWA and DYW2 to efficiently edit its target sites, clpP_69942 and rpoA_78691. To maximize the resemblance of this fusion with canonic DYW-PPRs, we aligned the C-termini of *Arabidopsis* E+-PPRs with DYW domains of DYW-PPRs (Fig. S1). Based on this alignment, we chose to remove the last 13 unconserved amino acids of CLB19 and the 3 first amino acids of the DYW domain of DYW2 (Fig. 2A). Noteworthily, this fusion reconstitutes the conserved ‘GY’ motif within the DYW domain (Fig. 1A, Fig. S1). The CLB19-DYW2 fusion construct was then transformed into the homozygous *nuwa-2* mutant containing the pABI3::*NUWA* construct (hereafter called *nuwa_ABI3_*) previously obtained to bypass the *nuwa* embryo lethality (30). Consistent with previously published results, the rpoA_78691 and clpP_69942 sites were 70-90% edited in Col-0 and strongly inhibited in *nuwa_ABI3_* plants (Fig. 2B) (30). Editing at both sites was partially restored to wild type levels in the *nuwa_ABI3_* lines complemented with CLB19-DYW2 (Fig. 2B). No increase in clpP_69942 and rpoA_78691 editing efficiency was observed in Col-0 plants expressing the CLB19-DYW2 fusion suggesting that the increase in editing efficiency was not due to the overexpression of the CLB19-DYW2 protein (Fig. 2B).

**FIGURE 2.**
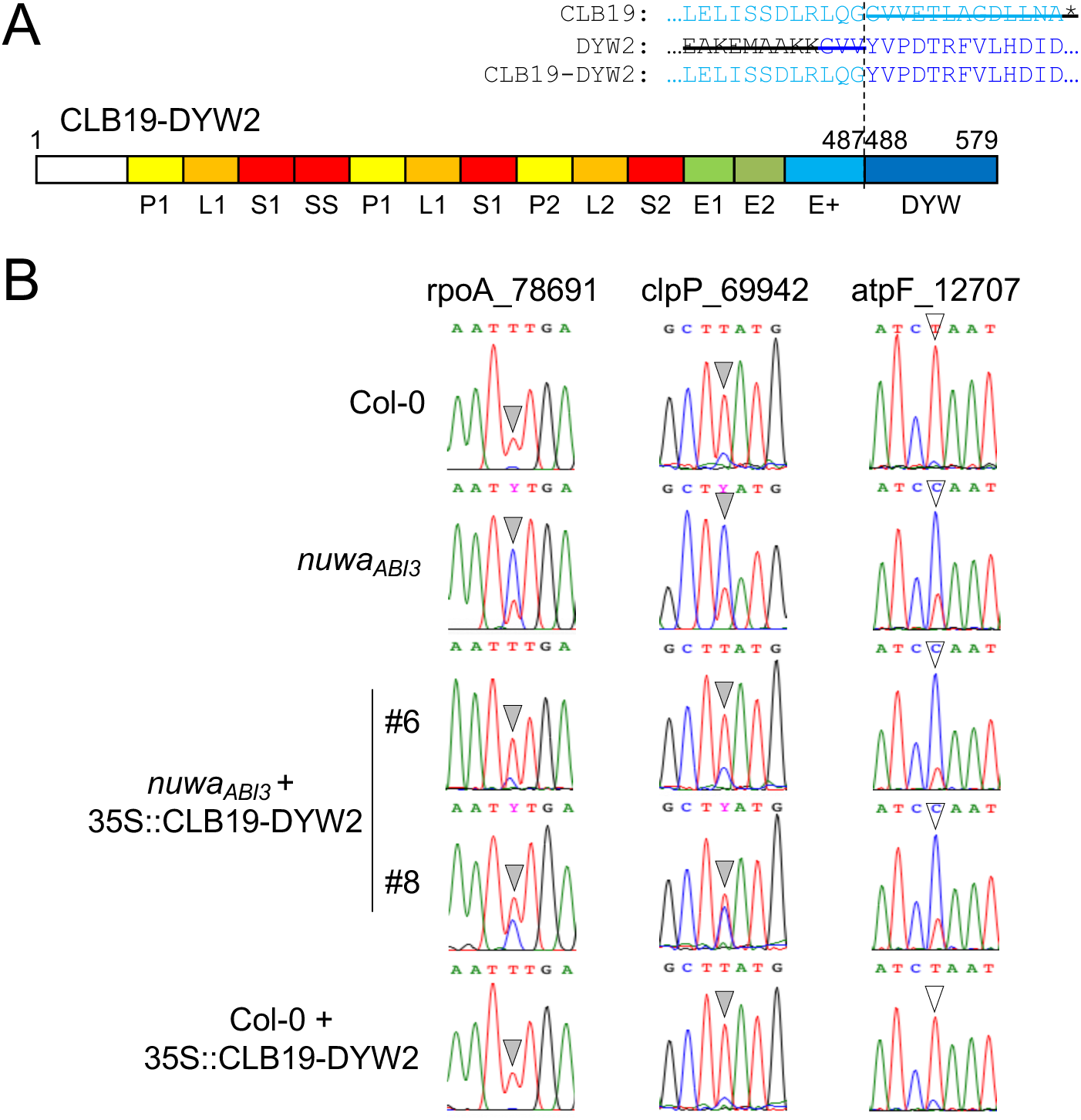
Complementation of the CLB19 editing sites by a CLB19-DYW2 fusion in the *nuwa-2* mutant. A) CLB19-DYW2 protein architecture: PPR tract, E, and E+ domains from CLB19 are fused to the DYW domain of DYW2. Original PPR amino acid numbers are indicated above their boundaries. Fusion area is detailed above the protein scheme. B) Sanger sequencing traces of plastid editing sites in Col-0, *nuwa_ABI3_* mutant, two independent T2 lines expressing the CLB19-DYW2 fusion in the *nuwa_ABI3_* background and one T2 line expressing the CLB19-DYW2 fusion in the Col-0 background. Grey arrowheads (left and middle columns) show rpoA_78691 and clpP_69942 editing sites (depending on CLB19), white arrowheads (right column) show atpF_12707 editing site (depending on AEF1).

To evaluate the specificity of the complementation, we also analyzed editing efficiency of the plastidial atpF_12707 site which is recognized by AEF1/MPR25, an E+-PPR (41) and is dependent on DYW2 and NUWA (30) (Fig. 2B). Editing efficiency of the atpF_12707 site in the lines expressing the CLB19-DYW2 fusion was similar to that observed in *nuwa_ABI3_* plants (Fig. 2B), confirming that the increase of editing efficiency previously observed was restricted to the CLB19 editing sites.

Taken together, our results show that the NUWA protein is no longer required for the proper function of CLB19 and DYW2 proteins when they are fused together into the same protein. More generally, they suggest that an editing site depending on NUWA could become independent once its specific PPR carries a DYW domain.

### Truncated versions of CRR28 are able to complement *crr28-2* mutant

One hypothesis to explain that *Arabidopsis* DYW-PPRs truncated of their own DYW domain (ΔDYW-PPRs) can complement their respective mutant is that they recruit in *trans* another DYW domain to perform deamination (17, 20, 22, 24, 25). We therefore decided to investigate this hypothesis using two truncated versions of the DYW-PPR CRR28 that is required to edit the ndhB_96698 and ndhD_116290 sites in plastid (24). The first truncation, called CRR28ΔE+DYW, was produced by removing the amino acids downstream the 27^th^ amino acid of the CRR28 E+ domain (Fig. 3A). This was designed to mimic an E2-PPR similar to the CRR4 PPR protein, the only E2-PPR in *Arabidopsis*, whose C-terminal region extends 27 amino acids downstream of the E2 domain (Fig. S2). For the second truncated CRR28, called CRR28ΔDYW, we removed the whole DYW domain (according to the old DYW domain definition (12), Fig. 1) (Fig. 3B) to mimic an E+-PPR. The same construct has previously been produced by Okuda *et al* in 2009 (24).

**FIGURE 3.**
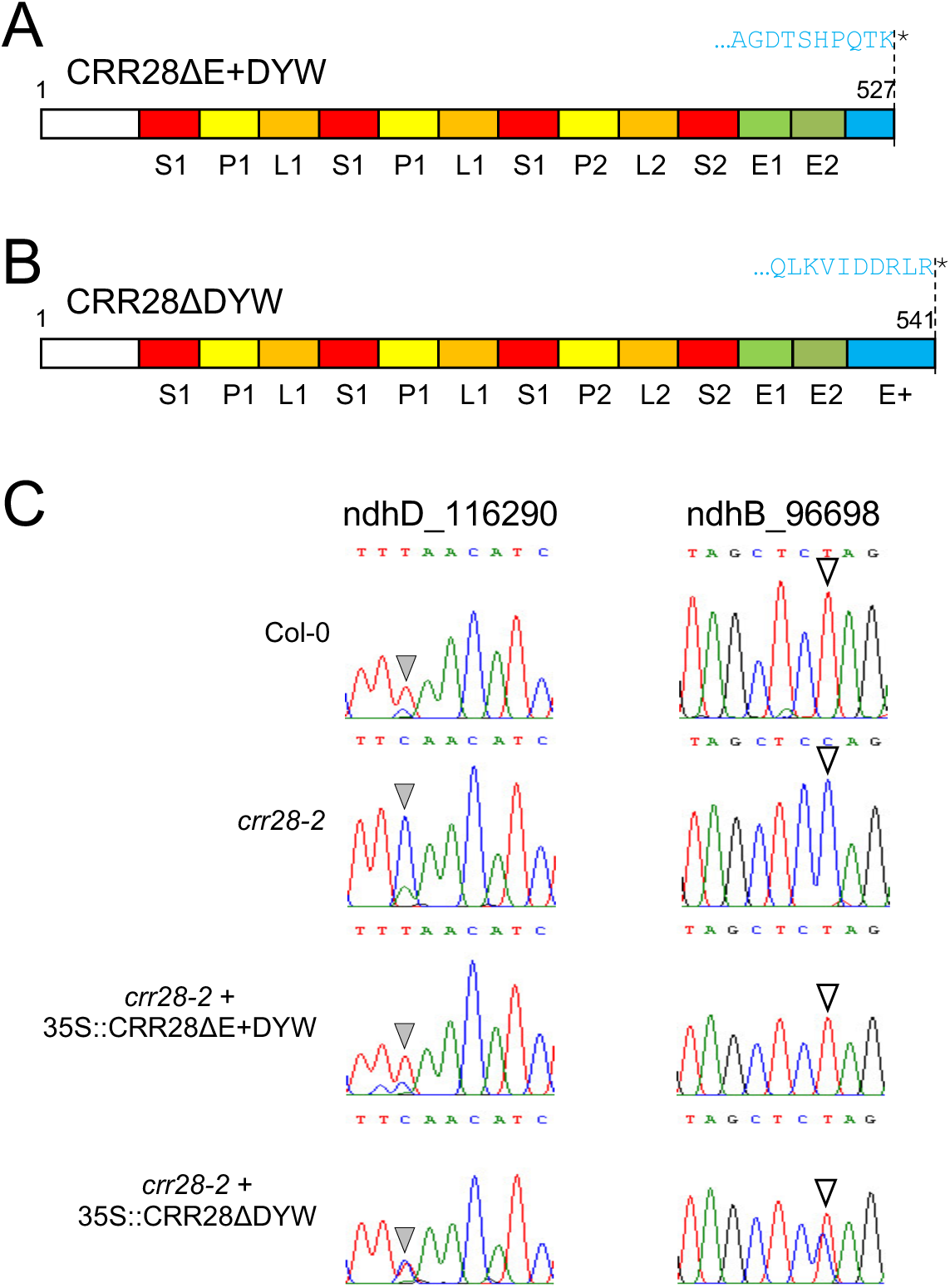
Complementation of the *crr28-2* mutant with truncated CRR28. A) CRR28ΔE+DYW protein architecture: CRR28 is truncated after the 27^th^ amino acid of its E+ domain. The last 10 amino acids of the truncated protein are indicated above the protein scheme. B) CRR28ΔDYW protein architecture: CRR28 is truncated at the end of the E+ domain. The last 10 amino acids of the truncated protein are indicated above the protein scheme. C) Sanger sequencing traces correspond to the CRR28 editing sites in Col-0, *crr28-2* mutant and *crr28-2* mutant expressing CRR28ΔE+DYW or CRR28ΔDYW constructs. Grey arrowheads (left column) show ndhD_116290 editing site, white arrowheads (right column) show ndhB_96698 editing site.

We then analyzed RNA editing efficiencies at the CRR28 editing sites in homozygous *crr28-2* mutant plants complemented by the truncated version of CRR28. As expected, editing at the two CRR28 sites was totally abolished in the mutant plants and restored to WT like levels in the CRR28ΔE+DYW complemented lines (Fig. 3C, Fig. S3A). RNA editing was also restored in the CRR28ΔDYW complemented lines albeit at a lower efficiency than in WT plants (Fig. 3C, Fig. S3B). It is worth noting that T1 plants showed a WT editing extent at ndhD_116290 site (Fig. S4). Altogether, our results show that CRR28 truncations mimicking E+-PPR or E2 (CRR4 like) C-terminus are able to functionally replace CRR28 protein in the *crr28-2* mutant.

**FIGURE 4.**
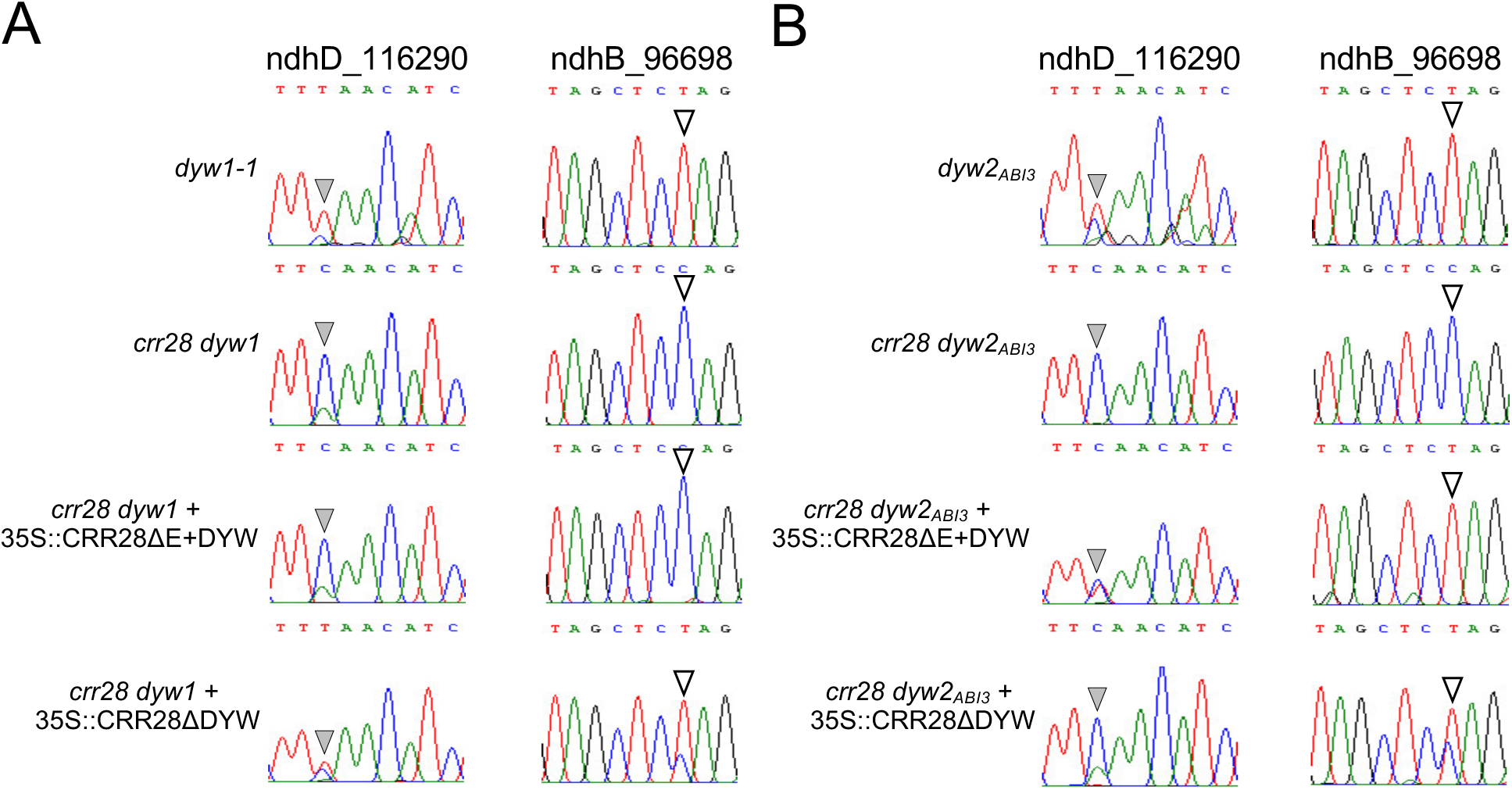
Editing efficiencies in plants expressing truncated CRR28 in absence of DYW1 or DYW2. Sanger sequencing traces correspond to the CRR28 editing sites in F3 plants obtained after crossing a *crr28-2* complemented lines with the *dyw1-1* (A) or the *dyw2_ABI3_* (B) mutants. Single *dyw1-1* or *dyw2_ABI3_* mutants are displayed on the first row, double mutants expressing no truncated PPR on the second row, double mutants expressing the CRR28ΔE+DYW constructs on the third row and finally double mutants expressing the CRR28ΔDYW constructs on the last row. Grey arrowheads (left columns) show ndhD_116290 editing site, white arrowheads (right columns) show ndhB_96698 editing site.

### Truncated CRR28 proteins require DYW1 or DYW2 to edit CRR28 sites

Our hypothesis was that the restoration of the editing defects in the *crr28-2* mutant by the truncated CRR28 was due to their interactions with DYW1-like proteins. More specifically, CRR28ΔDYW would require DYW2 because it is similar to an E+-PPR and CRR28ΔE+DYW would require DYW1 because it mimics CRR4, the only *Arabidopsis* plastidial E2-PPR. We therefore crossed complemented *crr28-2* plants expressing either CRR28ΔE+DYW or CRR28ΔDYW constructs (T1) with the homozygous *dyw1-1* mutant or the heterozygous *dyw2-1* mutant expressing the pABI3::*DYW2* construct, hereafter called *dyw2_ABI3_* (30). We then quantified RNA editing at the CRR28 sites in the F3 generation plants.

As expected, the *dyw1-1* single mutant showed a wild-type editing efficiency at CRR28 sites (Fig. 4A, Fig. S5C) whereas editing of CRR28 sites was completely abolished in the *crr28 dyw1* double mutant (Fig. 4A, Fig. S5E). The CRR28ΔE+DYW construct was not able to complement the editing defects of the CRR28 sites in the plants lacking both DYW1 and CRR28 (Fig. 4A, Fig. S3C). This indicates that the CRR28ΔE+DYW protein requires the presence of DYW1 to be functional. On the other hand, the CRR28ΔDYW construct displayed the same partial complementation in both the *crr28-2* and *crr28 dyw1* mutants (Fig. 4A, Fig. 3C, Fig. S3B, Fig. S3D), indicating that CRR28ΔDYW does not require the DYW1 protein to edit the CRR28 sites.

Similarly, the *dyw2_ABI3_* mutant edited the CRR28 sites whereas editing was abolished in the *crr28 dyw2_ABI3_* double mutant (Fig. 4B, Fig. S5D, Fig. S5F). In plants lacking both DYW2 and CRR28 and expressing the CRR28ΔDYW construct, editing defects were observed at both sites with a complete abolition of editing at the ndhD_116290 one (Fig. 4B, Fig. S3F). This indicates that CRR28ΔDYW, similarly to what was previously described for all E+-PPR, is dependent on DYW2. The *crr28 dyw2_ABI3_* lines expressing CRR28ΔE+DYW were, however, able to edit both editing sites at *dyw2_ABI3_* levels, demonstrating that CRR28ΔE+DYW does not require DYW2 for its editing activity (Fig. 4B, Fig. S3E, Fig. S5D).

In summary, our results show that in order to complement the *crr28-2* editing defects, the CRR28ΔE+DYW construct, mimicking the E2-PPR CRR4, requires the presence of DYW1 but does not depend on DYW2. In contrast, the CRR28ΔDYW construct, mimicking an E+-PPR, needs the presence of DYW2 to edit the CRR28 sites but does not require the presence of DYW1.

## DISCUSSION

### CRR28 deletions mimic PLS-DYW family evolution in Angiosperms

The origin and evolutionary history of DYW-PPR proteins is intimately linked to RNA editing (42). The association on a single protein of both a specific RNA binding domain (the PPR track) and a deamination domain (the DYW deaminase) allows the precise selection and deamination of the cytosines that require editing. As an illustration, all the editing sites in the moss *Physcomitrium patens* have been attributed to individual DYW PPR proteins (43). The situation is however more complex in vascular plants where the DYW domains of some editing factors have been shown to be dispensable for the deamination reaction. Additionally, several editing factors do not bear a DYW domain and belong to the E2- and E+-PPRs family, suggesting dissociation between the RNA binding and deamination activities necessary to the reaction. This idea is supported by the identification of the DYW1-like proteins that are short PPR proteins which only harbor the deaminase domain and can interact with non DYW PPR to perform RNA editing. This model is perfectly illustrated by the interaction between the E2-PPR CRR4 and DYW1 in *Arabidopsis* (28).

According to this hypothesis, the possibility of an interaction with DYW1-like proteins allowed the apparition of the E2- and E+-PPR subfamily in vascular plants, following the loss or degeneration of their DYW domain (then their E+) domains. In this model, a PPR harboring a DYW domain in an organism could lose it in another organism. Such a situation has been reported for CRR28: while it is a DYW-PPR protein in *Arabidopsis*, orthologs in *Lactuca sativa* and *Cynara cardunculus* are E+-PPR lacking a DYW domain (43). We therefore decided to put this model to test by artificially recreating CRR28 proteins devoid of their DYW domains and systematically test their dependency on DYW1-like proteins to perform editing. Our two deletions of the PPR-DYW CRR28 protein mimic the loss of domains in E+-PPRs and the plastidial E2-PPR CRR4 during evolution. They were introduced in the *crr28* mutants no longer expressing either DYW1 or DYW2 to test their requirement for these proteins (Fig. 3, Fig. 4).

### PPRs without DYW domain are depending on DYW1-like members

This approach led to the conclusion that CRR28ΔE+DYW harboring a “half” E+ at its C-terminus has exactly the same behavior as CRR4 and is depending on DYW1 (Fig. 4). Similarly, we showed that CRR28ΔDYW is behaving exactly as every other studied E+-PPRs and that its RNA editing activity relies on DYW2. Although E+-PPRs are all depending on DYW2, many E+-PPR editing sites were shown to be still partially edited in the *dyw2_ABI3_* mutant (30). Similarly, a single E+-PPR could impact its multiple target sites with different efficiency in the *dyw2_ABI3_* mutant (30, 35). This is exactly the behavior we observed in the *crr28 dyw2_ABI3_* + CRR28ΔDYW line, where editing at the ndhD site is completely abolished while the ndhB site is only partially affected (Fig. 4). Noticeably, two populations were observed among the F3 generation of *crr28 dyw2_ABI3_* + CRR28ΔE+DYW plants, one exhibiting better editing efficiencies than the other (Fig. S3E). Although the rationale behind these discrepancies is not clear, a plausible explanation could lie in the genetic complexity of our lines and in transgene silencing.

It is worth noting that CRR28ΔE+DYW is only 14 amino acids shorter than CRR28ΔDYW. It is therefore the absence of these 14 amino acids that determines the specific requirement of DYW1 versus DYW2. A simple explanation would be that the 14 amino acids determine the structural reconstitution of the DYW active site as suggested by Yang *et al* (2022) (36), an hypothesis that would require structural characterization of truncated PPRs interacting with DYW1 or DYW2 to be confirmed. Alternatively, scaffolding proteins like NUWA and GRP23 (see below) might also be involved in this specificity. The fact that we could identify orthologs of DYW2 (LsDYW2: XP_023754705.1; CcDYW2: XP_024959394.1) and NUWA (LsNUWA: XP_023756944.1; CcNUWA: XP_024970526.1) in *L. sativa* and *C. cardunculus,* the two plants that contain truncated versions of CRR28 (Fig. S6) is in agreement with this possibility.

### DYW domain specificity did not affect the complementation experiments

Several authors have recently described that the DYW domain of PPR proteins could be involved in the specificity of editing sites. This is the case, for example, in *Physcomitrium* where DYW domains are functionally different between PpPPR56 and other mitochondrial PPR editing factors, and in which residues 37–42 are involved in site-specific editing (14). Maeda *et al* (2022) using *E. coli* RNA editing system and *Arabidopsis*/*Physcomitrium* chimeric proteins, proposed that each DYW domain shows a distinct preference for neighboring nucleotides of the target site and thus participate in the editing specificity (44). DYW domains can also have a long-range impact on RNA recognition (45) and it was proposed that fine-tuning of the target specificity can be modulated by the DYW domain itself (46). This selectivity of some of the DYW domains could theoretically have complicated our domain complementation model and impaired our CRR28 protein deletion assays. Nevertheless, no such effect was observed in our experiments. In contrast, while DYW1 has been described to have a small preference for the ndhD_116290 site (UCA triplet) over the ndhB_96698 (CCA triplet)(44) a better editing efficiency was observed at the ndhB_96698 site in all our experiments (Fig. 3).

### NUWA, a protein scaffold stabilizing DYW2 and E+-PPR interactions

Pure PPR proteins like NUWA and GRP23 have been proposed to be scaffolding proteins in the editosome, helping to maintain the functional interactions between the partners (30, 31, 33, 36). In our experiment, the interaction between the truncated CRR28 and the DYW1-like proteins could for example be facilitated by these proteins. We tested this hypothesis using another plastidial E+-PPR, CLB19, to investigate whether NUWA is still required to edit E+-PPR sites when the E+-PPR protein is fused to a DYW domain. If the role of NUWA was to allow the interaction between the PPR protein bringing the RNA binding domain and the DYW1-like protein bringing the deaminase activity, it should therefore become dispensable when the two domains are carried by a single protein. In absence of NUWA, CLB19 and DYW2 are not able to efficiently edit CLB19 sites leading to a partial editing defect in the *nuwa_ABI3_* mutant. In contrast, in the same *nuwa_ABI3_* background, a fusion between CLB19 and the DYW domain of DYW2 is able to edit CLB19 sites, indicating that editing has become independent of the NUWA protein (Fig. 2). This result clearly supports the scaffold function of NUWA as a stabilizer of the E+-PPR/DYW2 interactions. Another PPR with a scaffolding function similar to NUWA has recently been described, indicating that this mechanism might be a general feature of the plant editosome (36).

### Toward a global model for the editosome core

PPR editing factors were first thought to be acting in a one PPR/one editing site fashion, without interacting with any other protein. Then, a more complex model for the editosome emerged (2, 3) in which the PPR tract of the PPR protein, helped by other proteins such as MORF/RIP proteins (47, 48), selectively binds the *cis* element surrounding the RNA editing site and the E+/DYW domain performs the cytidine-to-uracil editing reaction. Additionally, the E1/E2 domain is also involved in RNA binding (49), protein-protein interactions with other editosome components (50) and participates to the active site (23, 32).

Finally, seminal works on DYW1-like proteins (28–30, 33, 36, 39, 51) suggested that the functions carried by the PPR proteins could be split between several individual proteins that could then be assembled to recreate a true bona fide editing factor. Our results are clearly in favor of this ever more complex editosome in which highly specific interactions between a site-recognition PPR lacking the deamination C-terminal part and a short DYW1-like PPR would enable the structural and molecular reconstitution of the active site of full-length DYW-PPR. The appearance of the small DYW1-like subfamily in the vascular plants probably increased the resilience and flexibility of the molecular editing machinery by allowing the appearance and maintenance of C-terminal deletions that occurred during the gene rearrangements observed with the very rapid expansion of the family (52, 53).These interactions are then made possible by the action of scaffolding proteins whose precise mode of action still remain to be elucidated.

## EXPERIMENTAL PROCEDURES

### Plant material

T-DNA mutant *crr28-2* (SALK_115133) was previously described in Okuda *et al*, 2009 (24) and was ordered from the NASC (Nottingham Arabidopsis Stock Centre) (54). Tilling mutant *dyw1-1* (G262:) was previously described in Boussardon *et al* (2012) (28). Embryo complemented mutant *dyw2_ABI3_* and *nuwa_ABI3_* were generated and described in Guillaumot *et al*, 2017 (30), they correspond to *dyw2-1* (GK_332A07) + pABI3::*DYW2* mutant and *nuwa-2* (SAIL_784_A11) + pABI3::*NUWA* mutant, respectively. Plants were grown at 20°C constant temperature under long day conditions (16h light/day).

### PPR ORF cloning, plant transformation and crossing

Sequences coding for CRR28 truncated versions surrounded by Gateway *att*B recombination sites were synthesized by Twist BioScience company (San Francisco, US). CRR28ΔDYW corresponds to the CRR28 541 first amino acids until the end of the E+ domain and CRR28ΔE+DYW corresponds to the CRR28 527 first amino acids until the 27^th^ amino acid of the E+ domain. The CLB19-DYW2 fusion was constructed by combining PCR products corresponding to *CLB19* ORF (1-487, encoding the full-length protein without the last 13 amino acids of the E+ domain) and the sequence encoding the DYW domain of DYW2 (488-579, corresponding to the DYW domain without its 3 first amino acids). PPR ORFs were cloned into the pDONR207 vector using Gateway BP clonase enzyme (Invitrogen), then subcloned using Gateway LR clonase enzyme (Invitrogen) into the pGWB2 vector (55) that allows expression under the 35S promoter without any tag. Homozygous *nuwa_ABI3_* or *crr28-2* plants were transformed using C58C1 pMP90 *Agrobacterium tumefaciens* by floral dip (56) and selected on half MS (Duchefa MO0221.0050) media containing 50µg/mL of Kanamycin and 25µg/mL of Hygromycin B. Following transfer in soil, selected plants were genotyped and analyzed for their ability to restore editing defects. Complemented *crr28-2* T1 were crossed with *dyw1-1* or *dyw2_ABI3_* mutants and F2 and F3 generations were obtained in order to identify specific genotypes. Editing analysis was performed on F3 segregating plants. Primers used for the genotyping and cloning are listed in Table S1.

### RNA extraction and RNA editing analysis

Leaf total RNA was extracted using the NucleoZOL protocol (Macherey-Nagel) followed by RNA purification using Agencourt RNA Clean XP beads (Beckman-Coulter). cDNA was synthesized from 100 ng of total RNA using SuperScript II (Invitrogen). After RT-PCR with primers surrounding editing sites, products were sequenced by the Eurofins company. All primers used in this study are provided in the Table S1.

For the editing analysis of the CRR28 sites, editing was assayed in at least 3 plants of each genotype among the F3 generation. Only traces from the plants exhibiting the highest editing efficiencies are displayed in the Fig.3-4. All the other traces for every genotypes are displayed in supplemental figures (Fig. S3, Fig. S5).

## ACCESSION NUMBERS

*CLB19* (At1g05750), *CRR28* (At1g59720), *DYW1* (At1g47580), *DYW2* (At2g15690), *NUWA* (At3g49240)

## SUPPORTING INFORMATION

This article contains supporting information.

## AUTHOR CONTRIBUTIONS

C. L., K. B. and W. M. designed the research and supervised the experiments; K. B., D. M. and C. L. performed experiments; K. B., B. C., W. M. and C. L. wrote the manuscript.

## FIGURE LEGENDS

FIGURE S1. **Alignment of the amino acids of the E+ and DYW domains from Arabidopsis E+- and DYW-PPR proteins.** The alignment of the 58 E+ domains of *Arabidopsis* E+-PPRs with the 88 DYW domains of *Arabidopsis* DYW-PPRs was performed with Clustal Omega with default parameters. For space and display reasons, only four DYW-PPRs are shown at the top of the figure. Black arrowhead shows the end of the E+ domain defined in Lurin *et al* (2004). Red line at position 63 shows the chosen fusion point in the CLB19-DYW2 fusion.

FIGURE S2. **Alignment of the amino-acids occurring after the E2 domain in Arabidopsis E2- and E+-PPRs.** The alignment of the 46 E2- and 58 E+-PPRs from Arabidopsis was performed using Clustal Omega with default parameters. Only six E+-PPRs are shown at the top. Names of E2-PPRs known to be involved in RNA editing were added beside their AGI number. E2-PPR OTP70 (At4g25270) is absent from the alignment because it has no amino-acid after its annotated E2 domain. The black arrow highlights CRR4.

FIGURE S3. **Editing efficiencies of the CRR28 editing sites in mutant backgrounds expressing the two truncated CRR28 proteins.** Sanger sequencing traces correspond to CRR28 editing sites in lines expressing CRR28ΔE+DYW (A, C, E) or CRR28ΔDYW (B, D, F) in *crr28-2* simple mutant (A, B), *crr28 dyw1* (C, D) or *crr28 dyw2_ABI3_* (E, F) double mutants. Three plants are shown in A, B, C, D and F; seven plants are shown in E. Grey arrowheads (left columns) show ndhD_116290 editing site, white arrowheads (right columns) show ndhB_96698 editing site.

FIGURE S4. **Editing efficiencies of the ndhD_116290 editing site in two *crr28-2* T1 plants expressing the CRR28ΔDYW construct.** Sanger sequencing traces correspond to CRR28 ndhD_116290 editing site (shown by grey arrowheads) in two independent T1 plants.

FIGURE S5. **Editing efficiencies of the CRR28 editing sites in various mutant backgrounds.** Sanger sequencing traces correspond to CRR28 editing sites in Col-0 (A), *crr28-2* (B), *dyw1-1* (C), *dyw2_ABI3_* (D) simple mutants and in *crr28 dyw1* (E) and *crr28 dyw2_ABI3_* (F) double mutants. For each genotype, results obtained using three different plants are shown. Grey arrowheads (left columns) show ndhD_116290 editing site, white arrowheads (right columns) show ndhB_96698 editing site.

FIGURE S6. **Identification of putative DYW2 and NUWA orthologs in *Lactuca sativa* and *Cynara cardunculus*.** *Arabidopsis* protein sequences were used as BLASTP query on targeted organism. Best hit of both species were aligned with *Arabidopsis* proteins using Clustal Omega with default parameters. A) DYW2 alignment (AtDYW2: At2g15690; LsDYW2: XP_023754705.1; CcDYW2: XP_024959394.1). B) NUWA alignment (AtNUWA: At3g49240; LsNUWA: XP_023756944.1; CcNUWA: XP_024970526.1). PPR tracks, DYW domain and coiled-coil region in *Arabidopsis* proteins are indicated by yellow, blue, and green boxes above sequences, respectively.

