## Supplemental Table and Figures for "Dissecting the molecular puzzle of the editosome core in Arabidopsis organelles"

### Supporting information

| Primer name | Sequence, 5' to 3' | Comments |
|---|---|---|
| **Genotyping** | | |
| SK115133_LP | CGATGAGTAGTCTTGCTTGCC | WT crr28-2 genotyping |
| SK115133_RP | CTTCCTCAGCTCTTCCATGTG | WT crr28-2 genotyping |
| p35S-B | AGTGGAAAAGGAAGGTGGCT | CRR28 construct detection and crr8-2 insertion genotyping |
| At_CRR28_transg_R | CGAAACAAGACTTCTCTCAGGC | CRR28 construct detection and crr8-2 insertion genotyping |
| DYW1_genF | GGAAGCTGCTTTCTCCATGA | dyw1-1 genotyping and mutation sequencing |
| DYW1_genR | TGGAAGTTGCATAAATGGCTTAT | dyw1-1 genotyping |
| dyw2-1_LP | TCAAAATCCTCAACAAGGTGG | dyw2-1 genotyping (WT) |
| dyw2-1_RP | GCAATCGCTAACCTCTCACTG | dyw2-1 genotyping (WT and insertion) and pABI3::DYW2 construct detection |
| GABI_O8474 | ATAATAACGCTGCGGACATCTACATTTT | T-DNA Gabi (dyw2-1 insertion genotyping) |
| nuwa-2_LP | TGACTTGACCAGTTTGCTGTG | nuwa-2 genotyping (WT) |
| nuwa-2_RP | TAATGGGTACTGTGCTGGAGG | nuwa-2 genotyping (WT and insertion) |
| Otp100_Rev_center | TTCTCTCCAACAGCTTCCTCA | pABI3::NUWA construct detection |
| SALK_LBb1.3 | ATTTTGCCGATTTCGGAAC | T-DNA Salk (nuwa-2 insertion genotyping) |
| ABI3prom_F | TTATTGTTTCATTTCCACTTCAACG | pABI3 construct detection |
| **Cloning** | | |
| CLB19DYW2_for | GATTTGCGATTACAGGGCTATGTTCCTGACACACGG | CLB19DYW2 cloning |
| CLB19DYW2_rev | CCGTGTGTCAGGAACATAGCCCTGTAATCGCAAATC | CLB19DYW2 cloning |
| DYW2_stop | TCCACCTCCGGATCACCAGTAATCCCCGCAAGAAC | CLB19DYW2 cloning |
| CLB19_start | GGAGATAGAACCATGGGTCTCCTTCCCGTCG | CLB19DYW2 cloning |
| **Editing** | | |
| chloro 141 R | TTTATGAGGCACAAACGGGA | clpP_69942 editing site amplification |
| clpP_69942_R | TGAACCGCTACAAGATCAACA | clpP_69942 editing site amplification |
| clpP_69942_seq | TCTTGGAAGCGGAAGAATTAC | clpP_69942 editing site sequencing |
| atpF_12707_F | TTCGTTTACTTGGGTCACTGG | atpF_12707 editing site amplification |
| atpF_12707_R | TTCGCGGACTTGATTAATTG | atpF_12707 editing site amplification |
| Chloro25F | TCCGTTTCTACGTTACGCAAG | atpF_12707 editing site sequencing |
| At_rpoA_F | GGACGCTTTATTCTGTCTCCAC | rpoA_78691 editing site amplification |
| At_rpoA_R | TGGCATTTTCAACAGGCATGA | rpoA_78691 editing site amplification |
| At_rpoA_seq | GTAAAATAATGTCTCGAGCAG | rpoA_78691 editing site sequencing |
| ndhD_116281_116494_F | CATGTGGGGTGGAAAGAAAC | ndhD_116290 editing site amplification |
| ndhD_116281_116494_R | AAGTGCAGCGCCAATAAATC | ndhD_116290 editing site amplification and sequencing |
| ndhB_95608-96698_F | GAGGAATGTTTTTATGTGGTGCT | ndhB_96698 editing site amplification |
| ndhB_95608-96698_R | CCGATTTGACCTATGGACGA | ndhB_96698 editing site amplification |
| NdhBedIII | ATTTCTTGAAGCTCAATCTCTC | ndhB_96698 editing site sequencing |

TABLE S1. **Primers used in this study.**

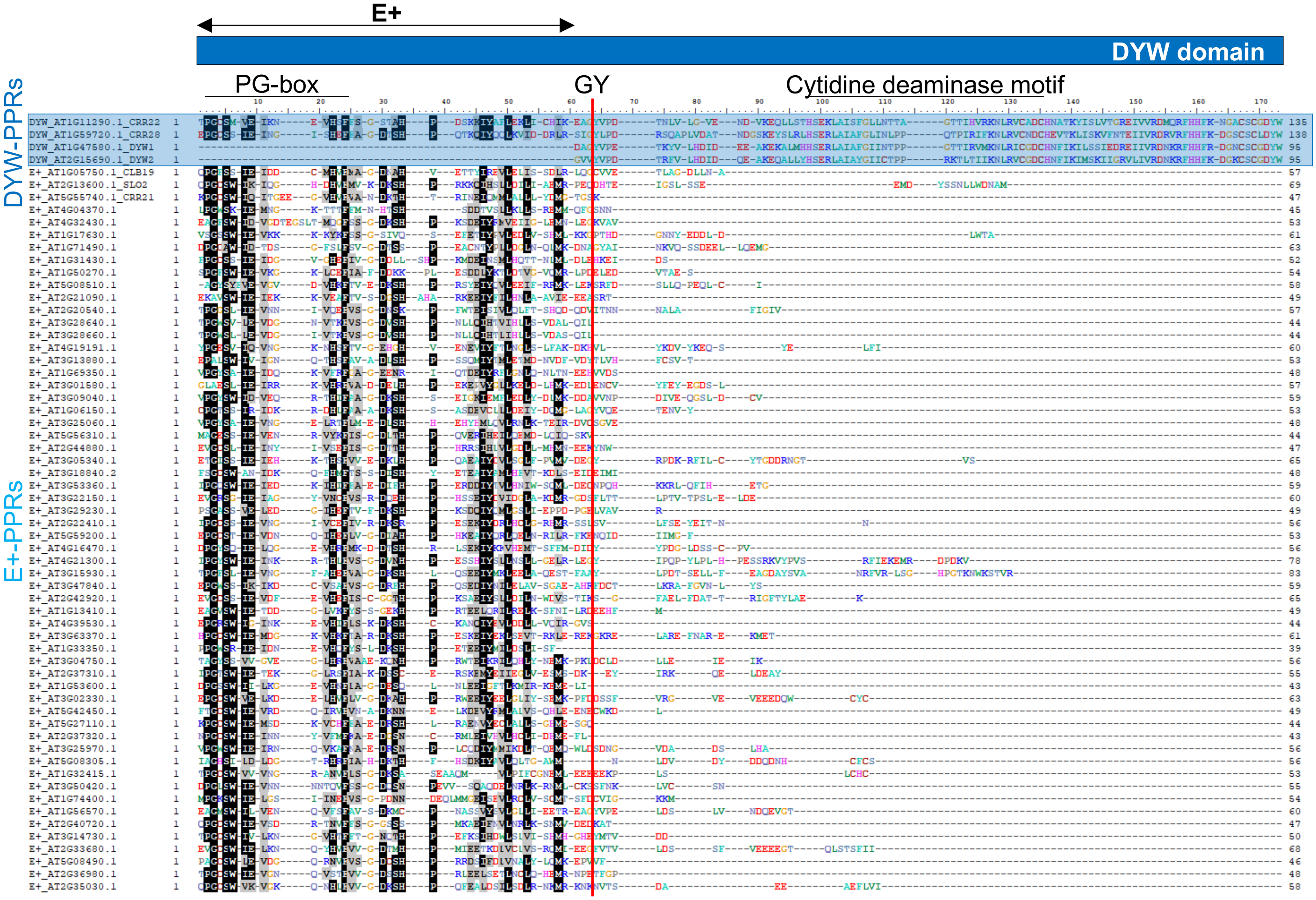


FIGURE S1. **Alignment of the amino acids of the E+ and DYW domains from Arabidopsis E+- and DYW-PPR proteins.**

The alignment of the 58 E+ domains of Arabidopsis E+-PPRs with the 88 DYW domains of Arabidopsis DYW-PPRs was performed with Clustal Omega with default parameters. For space and display reasons, only four DYW-PPRs are shown at the top of the figure. Black arrowhead shows the end of the E+ domain defined in Lurin et al (2004). Red line at position 63 shows the chosen fusion point in the CLB19-DYW2 fusion.

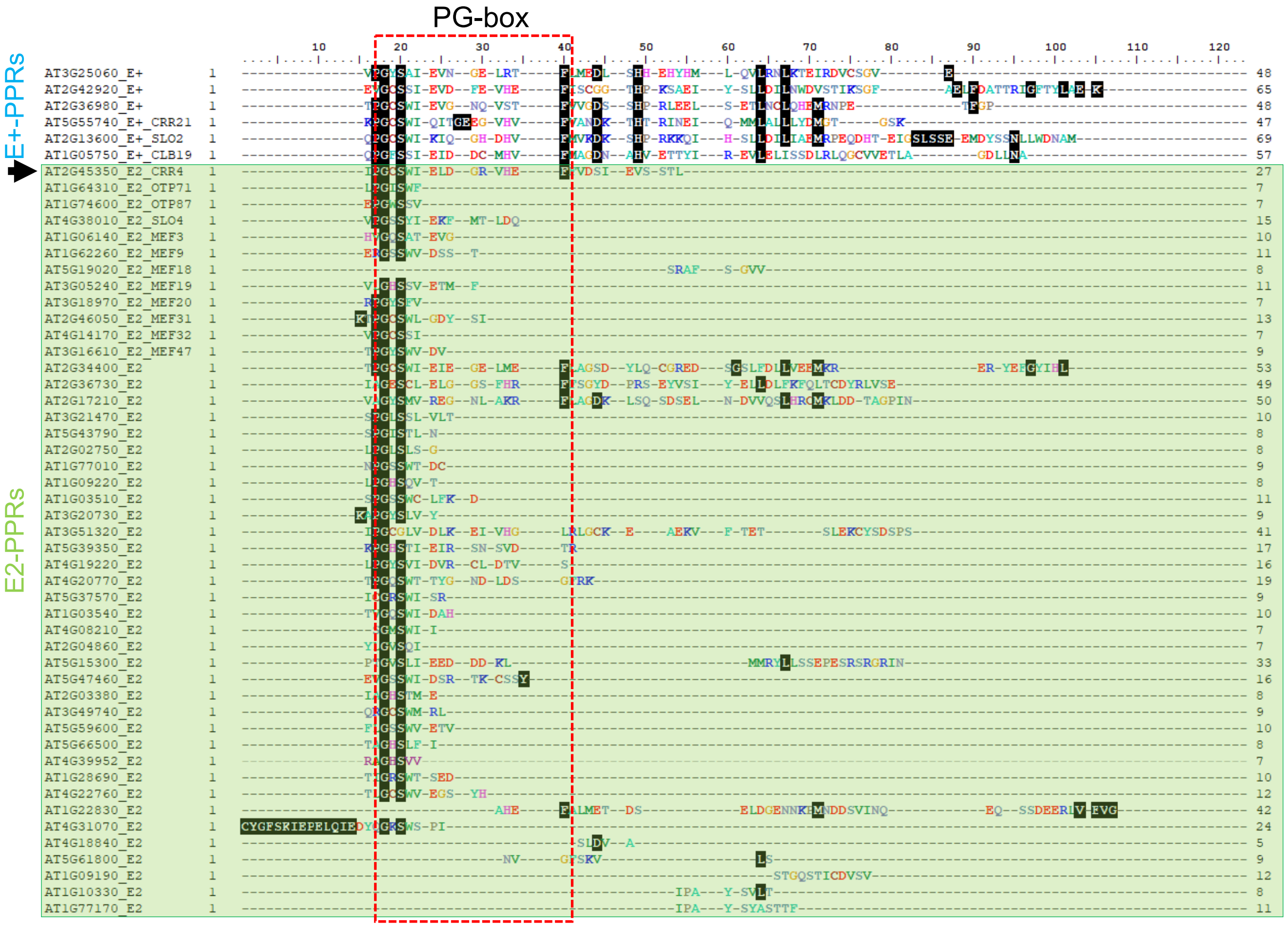


FIGURE S2. **Alignment of the amino-acids occurring after the E2 domain in Arabidopsis E2- and E+-PPRs.**

The alignment of the 46 E2- and 58 E+-PPRs from Arabidopsis was performed using Clustal Omega with default parameters. Only six E+-PPRs are shown at the top. Names of E2-PPRs known to be involved in RNA editing were added beside their AGI number. E2-PPR OTP70 (At4g25270) is absent from the alignment because it has no amino-acid after its annotated E2 domain. The black arrow highlights CRR4.

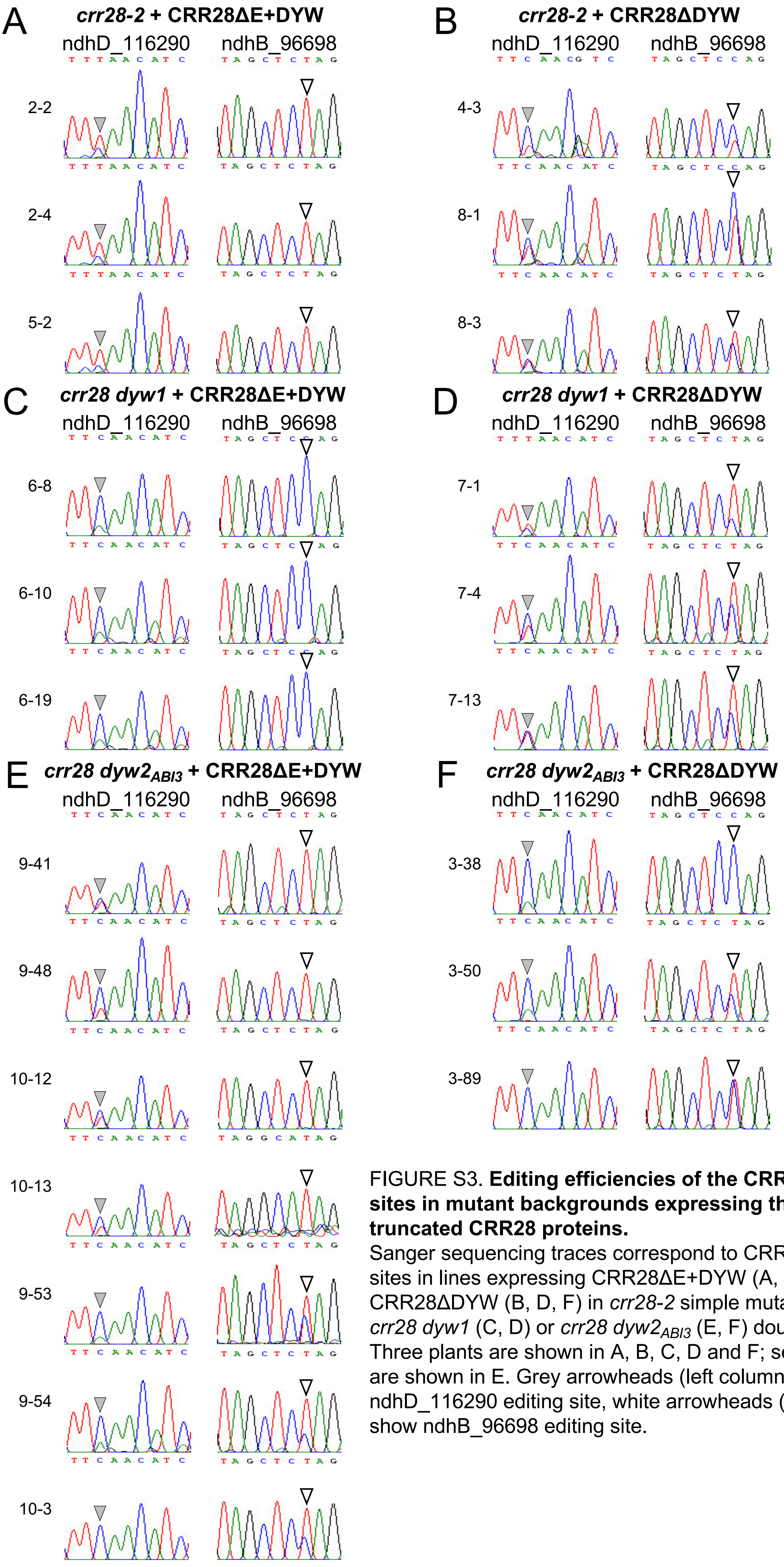


FIGURE S3. **Editing efficiencies of the CRR28 editing sites in mutant backgrounds expressing the two truncated CRR28 proteins.**
Sanger sequencing traces correspond to CRR28 editing sites in lines expressing CRR28ΔE+DYW (A, C, E) or CRR28ΔDYW (B, D, F) in *crr28-2* simple mutant (A, B), *crr28 dyw1* (C, D) or *crr28 dyw2*$_{ABI3}$ (E, F) double mutants. Three plants are shown in A, B, C, D and F; seven plants are shown in E. Grey arrowheads (left columns) show ndhD_116290 editing site, white arrowheads (right columns) show ndhB_96698 editing site.

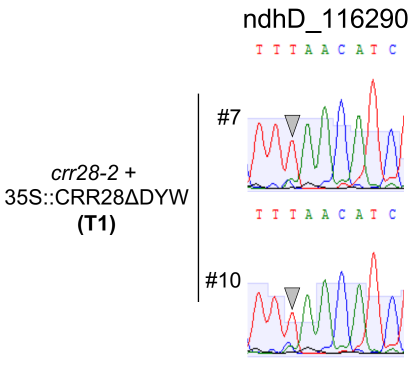


FIGURE S4. **Editing efficiencies of the ndhD_116290 editing site in two crr28-2 T1 plants expressing the CRR28ΔDYW construct.**
Sanger sequencing traces correspond to CRR28 ndhD_116290 editing site (shown by grey arrowheads) in two independent T1 plants.

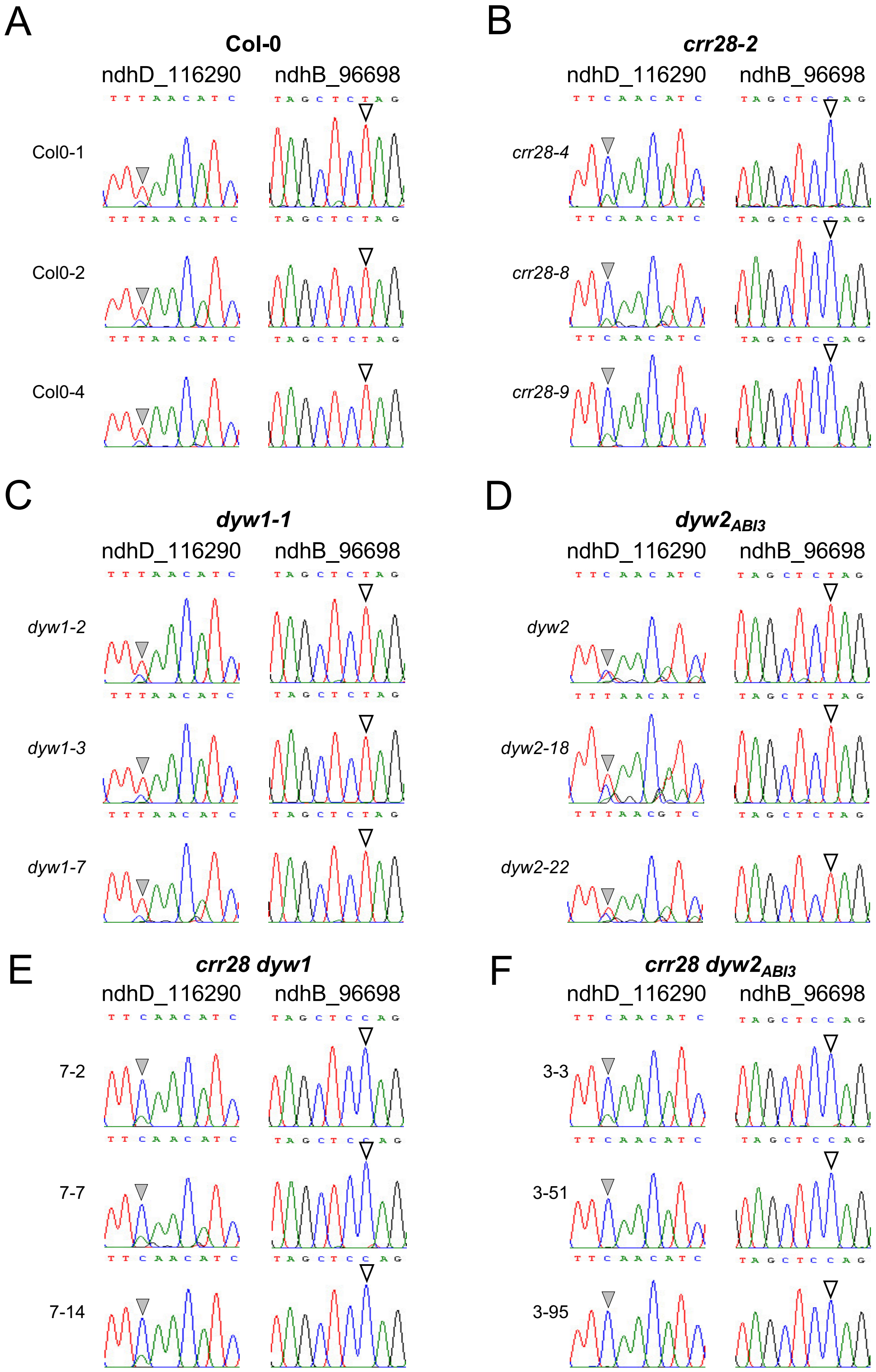


FIGURE S5. **Editing efficiencies of the CRR28 editing sites in various mutant backgrounds.**
Sanger sequencing traces correspond to CRR28 editing sites in Col-0 (A), *crr28-2* (B), *dyw1-1* (C), $dyw2_{ABI3}$ (D) simple mutants and in *crr28 dyw1* (E) and *crr28* $dyw2_{ABI3}$ (F) double mutants. For each genotype, results obtained using three different plants are shown. Grey arrowheads (left columns) show ndhD_116290 editing site, white arrowheads (right columns) show ndhB_96698 editing site.

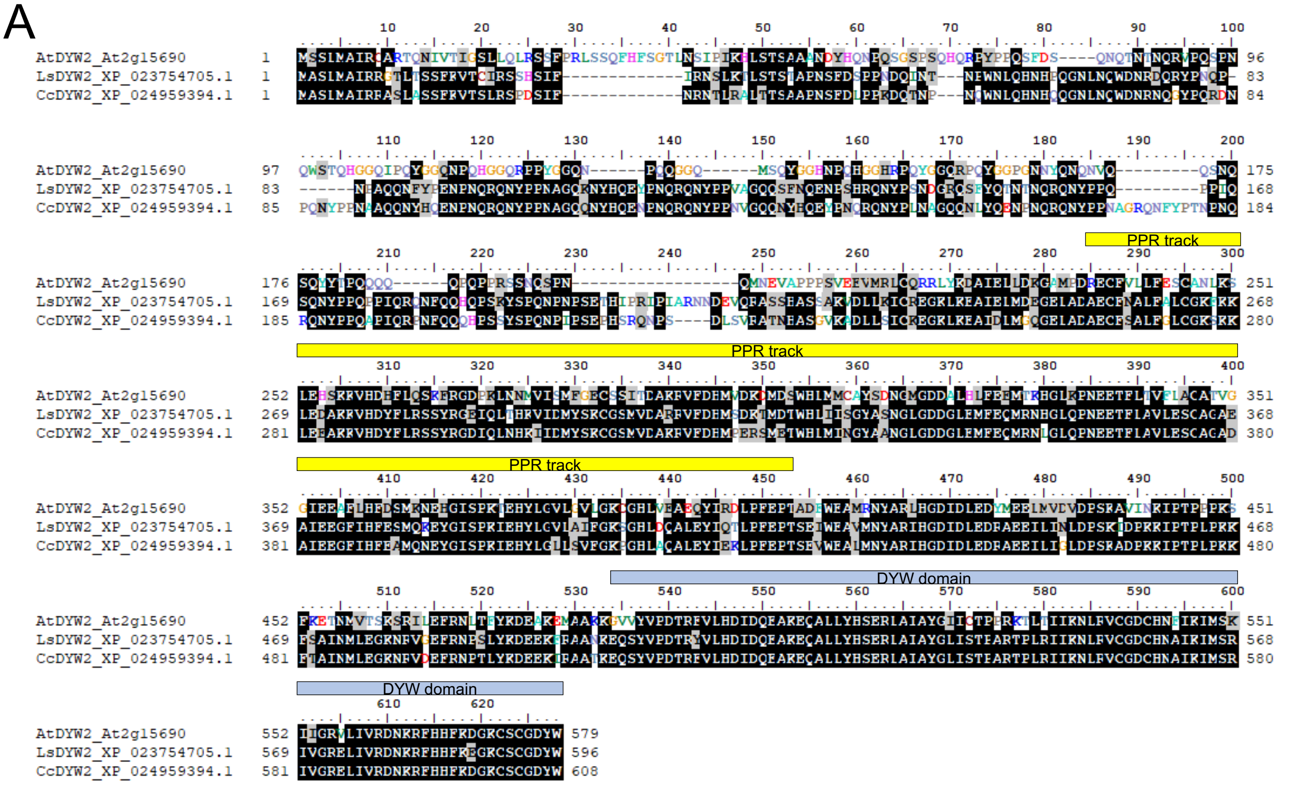

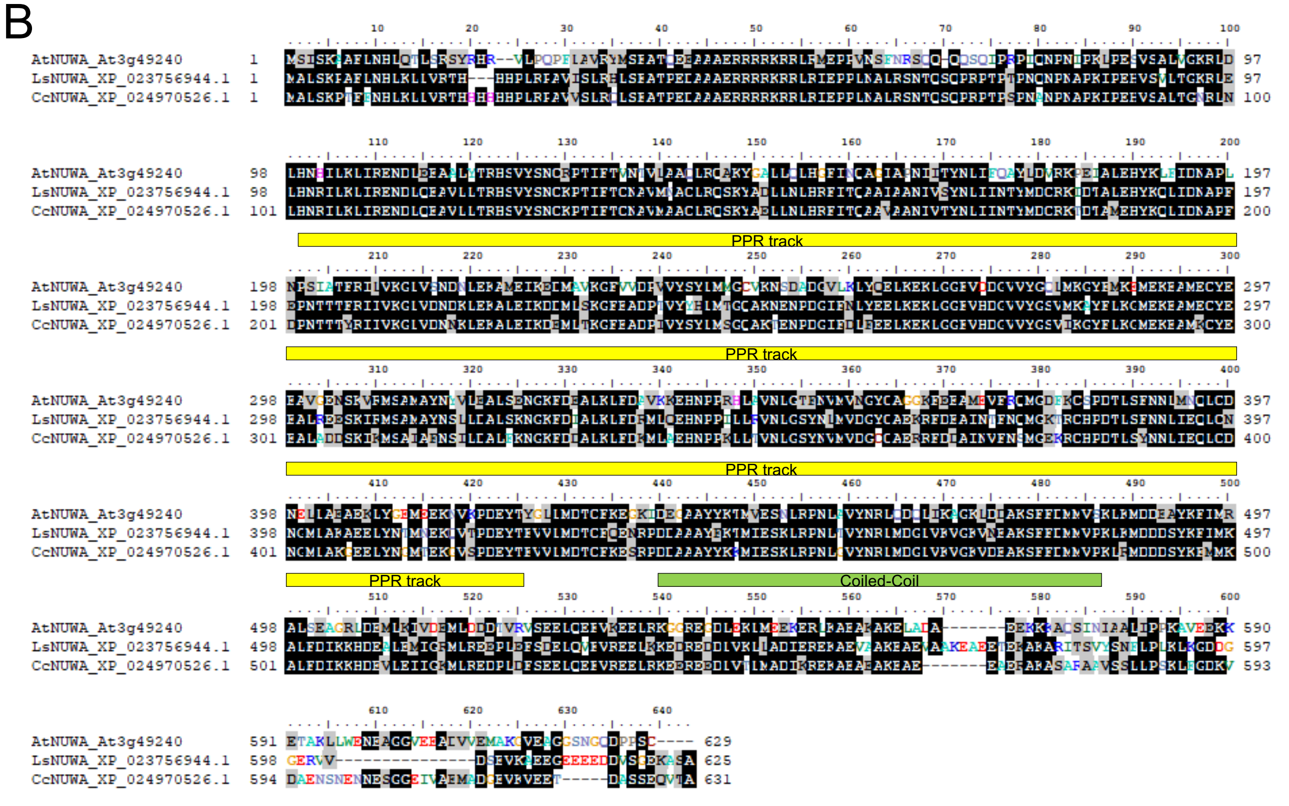

FIGURE S6. **Identification of putative DYW2 and NUWA orthologs in *Lactuca sativa* and *Cynara cardunculus*.**
Arabidopsis protein sequences were used as BLASTP query on targeted organism. Best hit of both species were aligned with Arabidopsis proteins using Clustal Omega with default parameters. A) DYW2 alignment (AtDYW2: At2g15690; LsDYW2: XP_023754705.1; CcDYW2: XP_024959394.1). B) NUWA alignment (AtNUWA: At3g49240; LsNUWA: XP_023756944.1; CcNUWA: XP_024970526.1). PPR tracks, DYW domain and coiled-coil region in Arabidopsis proteins are indicated by yellow, blue, and green boxes above sequences, respectively.
